# Comparative Evaluation of cobas Influenza A/B & RSV and Simplexa Flu A/B & RSV for Rapid Detection of Influenza and Respiratory Syncytial Viruses

**DOI:** 10.1101/2020.04.15.044099

**Authors:** Kyu-Hwa Hur, Heungsup Sung, Mi-Na Kim

## Abstract

We compared two molecular point-of-care tests, the cobas Influenza A/B & RSV (cobas Liat) and the Simplexa Flu A/B & RSV (Simplexa). A total of 236 respiratory specimens from patients referred for respiratory viruses testing were retrospectively evaluated; 53 specimens tested positive for each of Flu A, Flu B, and RSV, and 77 specimens tested negative based on the results of the reference method, i.e. the Seegene Allplex Respiratory Panel 1/2/3 (Seegene, Seoul, Korea). The turnaround time (TAT) was 20 min per specimen for cobas Liat and 78 min per eight specimens for Simplexa. The numbers of invalid results were one (0.4%) in cobas Liat and 10 (4.2%) in Simplexa (p < 0.05). All results were consistent with those of the reference method in cobas Liat. The sensitivity and specificity for Flu A, Flu B and RSV A were 100% with Simplexa. However, the sensitivity for RSV B was 80.0% with Simplexa, which was significantly lower than that of cobas Liat (p < 0.05). Comparison of the cycle threshold (Ct) values for RSV with Simplexa and the reference method showed the correlation as continuous variables (p < 0.001) with a higher propensity for obtaining Ct values with Simplexa, the exception being the six false negative results; their Ct values were more than 30 in the reference method. Cobas Liat showed high sensitivity for the detection of RSV B with rapid TAT, and a good workflow efficiency.

## INTRODUCTION

Respiratory virus infection is mostly associated with morbidity and mortality in all ages worldwide. It is also associated with increasing hospitalization costs and imposes an economic burden on the healthcare system (1, 2). Among the respiratory viruses, influenza viruses (Flu) and respiratory syncytial virus (RSV) are the most significant causes of morbidity and mortality during the period when respiratory viruses are abundant. These are especially fatal in children, the elderly, and in patients with comorbidities such as immunosuppression (2-5). Rapid and accurate diagnosis of Flu and RSV is essential to provide patients with quick and appropriate treatment, and reduce unnecessary healthcare-related expenses (6-8).

There are several laboratory methods for testing Flu and RSV, but rapid antigen detection tests are the most commonly used due to their ease of use and rapid turnaround time (TAT). However, they demonstrate a critical problem, i.e. poor sensitivity (8-10). Reverse transcription polymerase chain reaction (RT-PCR) has enabled the development of molecular diagnostic tests for the detection of Flu and RSV, which have become the gold standard test methods due to improved sensitivity and short TAT (6, 10). The sample-in, answer-out system is a fully automated test method, which is suitable as point-of-care (POC) test, and is easier to use, has short TAT, and demonstrates high performance.

Several automated multiplex RT-PCR tests for Flu and RSV have been approved by the Food and Drug Administration, and are currently available. In this study, we compared the cobas Influenza A/B & RSV (cobas Liat; Roche Diagnostics, Indianapolis, IN) with the Simplexa Flu A/B & RSV (Simplexa; Diasorin Molecular, Cypress, CA), and evaluated their performance against the routine RT-PCR method.

## MATERIALS AND METHODS

### Clinical specimens

A total of 236 retrospective respiratory specimens from patients who referred for respiratory viruses testing due to suspected viral respiratory infections were used in the study, of which 53 specimens tested positive for each of Flu A, Flu B, and RSV, and 77 specimens tested negative based on the reference method. The specimens were collected in 3 ml of universal viral transport medium (UTM, Copan Diagnostics Inc., Murrieta, CA) during the respiratory virus season of 2018 and 2019, and were stored at −80°C after routine clinical tests. The types of respiratory specimen included 205 nasopharyngeal (NP) swabs, 19 NP aspirates, and 12 bronchoalveolar lavage (BAL) fluids. The entire study was conducted at the Asan Medical Center (Seoul, Korea) by two expert laboratory technologists. All tests were performed as per the manufacturer’s instructions. This study was approved as “exempt review category” by the Institutional Review Board of Asan Medical Center due to the retrospective nature of the study and minimal risk to the patients.

### Study design

All specimens were tested concurrently using two cobas Liat Analyzer (Roche Diagnostics) instruments and a LIAISON MDX (Diasorin Molecular) instrument. For cobas Liat testing, 200 μl of the specimen was dispensed in the manufacturer’s cartridge using the supplied assay pipette and loaded onto the cobas Liat Analyzer. For Simplexa testing, 50 μl each of the reaction mix and specimen were dispensed into the wells of the direct amplification disc while being careful not to create bubbles and loaded onto the LIAISON MDX. The total hands-on time for starting the test was less than one minute per one specimen in cobas Liat and about 10 min per eight specimens in Simplexa. The tests took 20 min per one specimen in cobas Liat and 78 min per one test, which could process up to eight specimens in Simplexa. The Seegene Allplex Respiratory Panel 1/2/3 (Seegene, Seoul, Korea) was used as the reference method for Flu and RSV. Test operators were blinded to the reference results. When there was an amplification failure of internal control to the initial test, the result was considered invalid. Tests were repeated once on the respective instruments for the invalid results in the initial tests.

### Data analysis

The sensitivity, specificity, and 95% confidence intervals (CIs) of cobas Liat and Simplexa were determined in comparison to those of the reference method. The significant differences in both tests were directly compared using the McNemar test. As cobas Liat does not provide cycle threshold (Ct) values, comparison of the Ct values with the reference method was performed only with Simplexa using a simple linear regression analysis. All statistical analyses were performed using IBM SPSS Statistics 22.0 software (IBM Corp. 2013. Armonk, NY).

## RESULTS

The numbers of invalid results were one (0.4%; 1 NP swab) in cobas Liat and 10 (4.2%; 2 NP swabs, 4 NP aspirates, and 4 BAL fluids) in Simplexa (p < 0.05). All invalid results were retested with the same tests. The one invalid result was determined to be a valid result, likewise the reference method result, after retesting in cobas Liat, whereas in Simplexa, only two invalid results (1 NP swab and 1 BAL fluid) turned out to be valid results; eight invalid results remained invalid after retesting.

The results of Flu A, Flu B, RSV A, and RSV B for cobas Liat and Simplexa were compared with the reference results (Table 1). Initial invalid results for both tests were excluded in the analysis. There was no false positive and negative result in cobas Liat; the overall sensitivity and specificity for both Flu and RSV were 100%, whereas there were six false negative results for RSV B in Simplexa, and the sensitivity for RSV B was 80.0% (95% CI, 61.4– 92.3%). Of the six false negative specimens, four specimens were NP swabs and two specimens were NP aspirates. The sensitivity and specificity of Simplexa for Flu A, Flu B, and RSV A were 100%. The difference in the sensitivities of cobas Liat and Simplexa for RSV B was statistically significant (difference: 20.0%; 95% CI: 5.7–34.3%; p < 0.05).

**TABLE 1.**
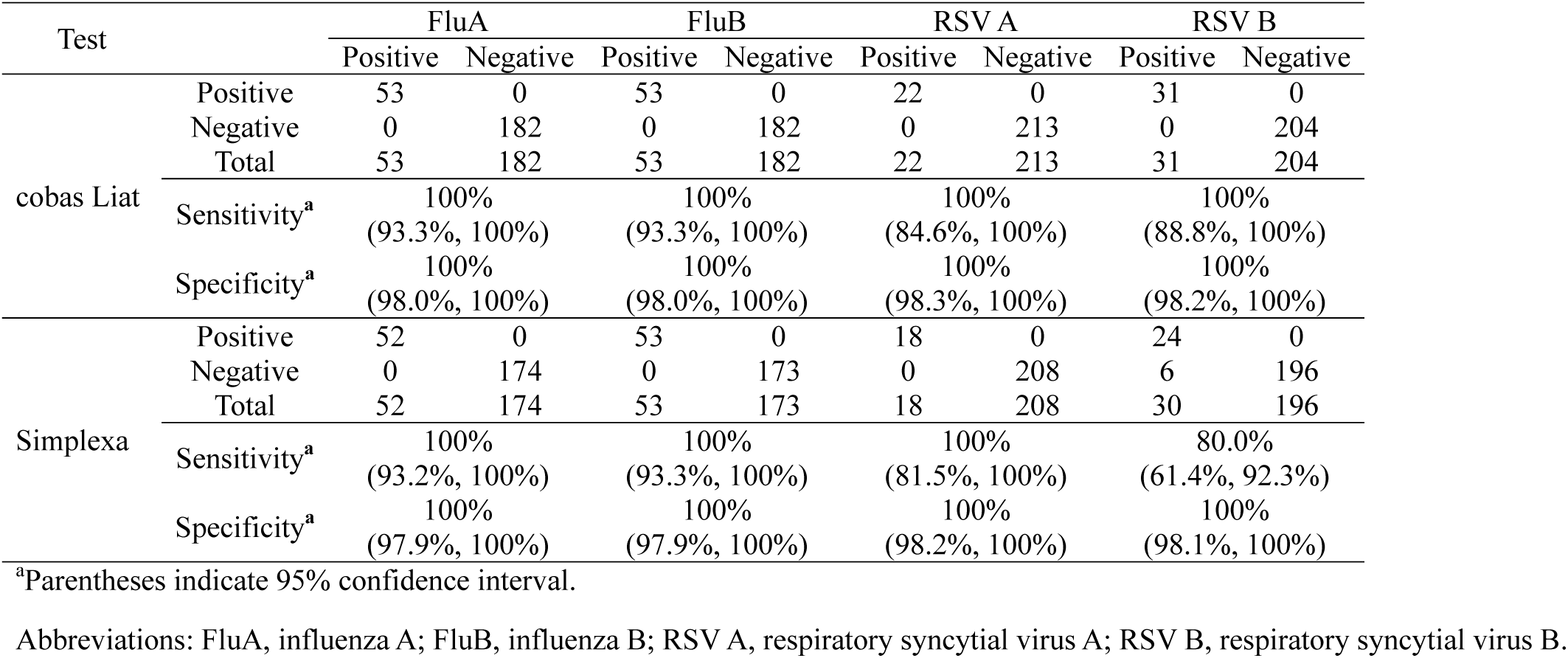
Performance of cobas Liat and Simplexa with respect to the detection of influenza A, influenza B, respiratory syncytial virus A and respiratory syncytial virus B

**TABLE 2.** Comparison of the two automated multiplex RT-PCR test methods with respect to test instruments, loading capacities, processing times, specimen volumes, and required materials

| Test kit | Test instrument | Loading capacity/test | Processing time (min) |  | Specimen volume (µl) | Required materials |
| --- | --- | --- | --- | --- | --- | --- |
|  |  |  | Hands-on time | Turnaround time |  |  |
| cobas Influenza A/B & RSV | cobas Liat Analyzer | 1 | <1 | 20 | 200 | None |
| Simplexa Flu A/B & RSV | LIAISON MDX | 1–8 | 1–10 according to the sample size | 78 | 50 | Reaction mix<br>Pipetting tools |

The Ct values of RSV for the reference method were compared with those in Simplexa (Fig. 1). Except for the six false negative results in Simplexa, we found the correlation between the Ct values as continuous variables (r2=0.305, p < 0.001) with a higher chance of obtaining Ct values with Simplexa. All six false negative results in Simplexa showed that the Ct values were >30 in the reference method.

**Figure 1.**
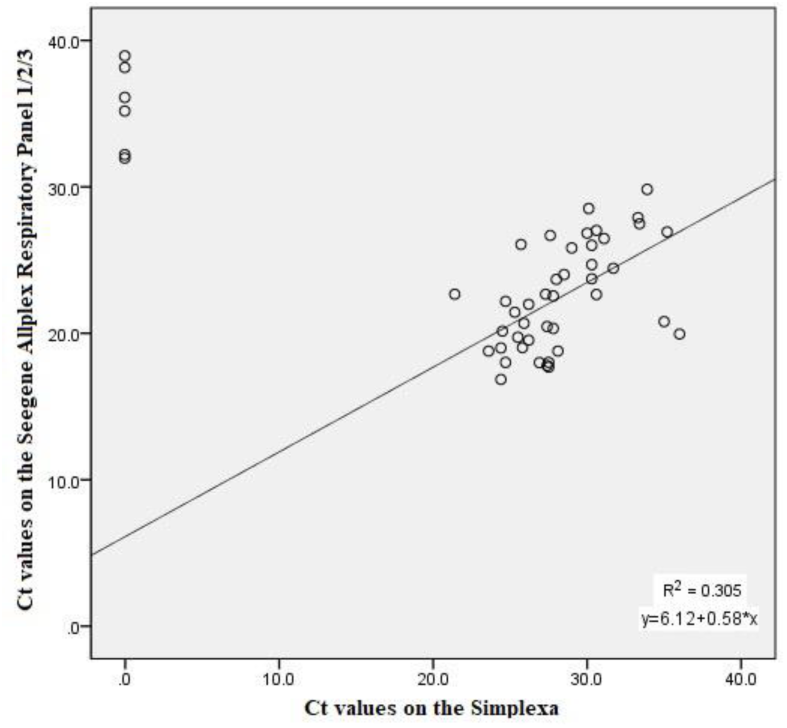
Scatter plot and simple linear regression analysis of the relationship between the Ct values of RSV on using the Seegene Allplex Respiratory Panel 1/2/3 (reference method) and Simplexa. Ct, cycle threshold

## DISCUSSION

The rate of invalid results for Simplexa was significantly higher than that for cobas Liat. If the rate of invalid results increases, there would be unnecessary wastage of resources and the workload of the laboratories would increase because of retesting. There can be several reasons for the increase in the rate of invalid results, including failure in the test procedure (11). For example, if bubbles are created when dispensing the specimen into the wells of the disc, the test is more likely to fail in Simplexa. Meanwhile, cobas Liat purifies extracted nucleic acids before the initiation of testing, while Simplexa does not include the purifying process. It is possible that the higher rate of invalid results in Simplexa may be due to the decrease in purity during the automated nucleic acid extraction process.

The sensitivity and specificity of both tests in this study are comparable to the results reported in previous studies (11-13). In our study, significant difference in the sensitivity for RSV B detection was demonstrated between Simplexa than of cobas Liat, with Simplexa showing a lower sensitivity than cobas Liat, and all six specimens of RSV B, which tested false negative in Simplexa, showed that the Ct values were more than 30 in the reference method. Comparing RSV A with RSV B, the lower sensitivity for RSV B detection may have resulted from the higher limit of detection for RSV B (14, 15) or due to the influence of sequence variants and probe-target mismatch (16). Given that the volume of specimen required for testing varies for different methods, this may explain the differences in test performance, rate of invalid results, sensitivity, and so on. The low volume of specimen needed for Simplexa makes it an ideal choice in cases where adequate amount of specimen is not available or where retesting is required; however, this may result in low sensitivity and high rate of invalid results. The additional dilution may also contribute to the poor sensitivity and high rate of invalid results. No dilution medium is added prior to testing in cobas Liat, while 50 μl of reaction mix is dispensed with the specimen in Simplexa.

Cobas Liat is known to have the shortest TAT per test among the real-time PCR-based molecular POC tests currently in use. Given that rapid detection is of utmost importance for the treatment of Flu and RSV, the use of cobas Liat is highly advantageous as a POC test. In addition, it is easy to use due to the fewer steps involved, and processes one specimen at a time, which reduces the risk of cross-contamination. Simplexa has a long unattended time, which makes it easier for the operator to perform other tasks. However, it is more cumbersome to process than cobas Liat, as it requires pipetting twice for every specimen and reaction mix and working carefully to prevent invalid or false results. The size of the analytical instrument is larger in Simplexa, and it requires a connection to the computer, while cobas Liat is smaller and can be operated only with its own instrument. Cobas Liat has the disadvantage of processing only three specimens every one hour. This is an important point where cobas Liat differs from the other POC tests, which means that multiple analyzers would be required as the number of specimens processed in the laboratory increases. This random-access method enables testing of individual specimens at a rapid TAT and increases laboratory workflow efficiency; however, it costs a lot to process large specimen volumes at once. In contrast, LIAISON MDX can process eight specimens in 78 min on the assumption of loading Simplexa to its full capacity. This batch-testing method is more useful for use in laboratories that process high specimen volumes (or during the influenza virus season); however, it is difficult to report results rapidly, which reduces the efficacy of this POC test. Cobas Liat is Clinical Laboratory Improvement Amendments (CLIA) waived, whereas Simplexa has moderate complexity for use with NP swab eluted in UTM. In this study, 19 nasal aspirates and 12 BAL fluids were analyzed using both tests. While Simplexa showed eight invalid and two false negative results, cobas Liat exhibited no invalid or false negative results for nasal aspirates and BAL fluids. It suggests that in addition to NP swabs, cobas Liat can process additional types of specimens. Additionally, the rate of invalid results in nasal aspirates and BAL fluids tended to be higher than that in NP swabs. This was the first attempt to evaluate POC tests with different types of specimens other than NP swabs. Although two previous studies have reported the results of tests using the same types of specimens as used in this study, they were not adequate for meaningful analysis as the number of specimens involved was too small (17, 18).

A limitation of our study is that the study was conducted with respiratory specimens stored at −80°C. Further, to prove that Simplexa cannot detect low positive specimens of RSV B, it is necessary to conduct further assessments with fresh specimens to obtain more precise results. The number of specimens was small, yet enough to demonstrate significant differences in the performances of the two POC tests. Cobas Liat seems to be a useful POC test for Flu and RSV compared with other real-time PCR-based molecular tests currently in use (11, 17-20).

In conclusion, the selection of a POC test method depends on the volume of samples, TAT, and laboratory resources, such as space, labor force, and capital. Upon considering the performance alone, cobas Liat can be used to provide efficient management of Flu and RSV in a POC setting.

## ACKNOWLEDGEMENTS

This study was supported by National Research Foundation of Korea grant funded by the Korea government (NRF-2016M3A9B6918716).

## REFERENCES

1. Molinari NA, Ortega-Sanchez IR, Messonnier ML, Thompson WW, Wortley PM, Weintraub E, Bridges CB. 2007. The annual impact of seasonal influenza in the US: measuring disease burden and costs. Vaccine 25: 5086–5096.

2. Thompson WW, Shay DK, Weintraub E, Brammer L, Cox N, Anderson LJ, Fukuda K. 2003. Mortality associated with influenza and respiratory syncytial virus in the United States. JAMA 289: 179–186.

3. Falsey AR, Hennessey PA, Formica MA, Cox C, Walsh EE. 2005. Respiratory syncytial virus infection in elderly and high-risk adults. New Engl J Med 352: 1749–1759.

4. Hall CB, Weinberg GA, Iwane MK, Blumkin AK, Edwards KM, Staat MA, Auinger P, Griffin MR, Poehling KA, Erdman D, Grijalva CG, Zhu Y, Szilagyi P. 2009. The burden of respiratory syncytial virus infection in young children. New Engl J Med 360: 588–598.

5. Nair H, Brooks WA, Katz M, Roca A, Berkley JA, Madhi SA, Simmerman JM, Gordon A, Sato M, Howie S, Krishnan A, Ope M, Lindblade KA, Carosone-Link P, Lucero M, Ochieng W, Kamimoto L, Dueger E, Bhat N, Vong S, Theodoratou E, Chittaganpitch M, Chimah O, Balmaseda A, Buchy P, Harris E, Evans V, Katayose M, Gaur B, O’Callaghan-Gordo C, Goswami D, Arvelo W, Venter M, Briese T, Tokarz R, Widdowson M-A, Mounts AW, Breiman RF, Feikin DR, Klugman KP, Olsen SJ, Gessner BD, Wright PF, Rudan I, Broor S, Simões EAF, Campbell H. 2011. Global burden of respiratory infections due to seasonal influenza in young children: a systematic review and meta-analysis. Lancet 378: 1917–1930.

6. Rogan DT, Kochar MS, Yang S, Quinn JV. 2017. Impact of Rapid molecular respiratory virus testing on real-time decision making in a pediatric emergency department. J Mol Diagn 19: 460–467.

7. Rolfes MA, Yousey-Hindes KM, Meek JI, Fry AM, Chaves SS. 2016. Respiratory viral testing and influenza antiviral prescriptions during hospitalization for acute respiratory illnesses. Open Forum Infect Dis 3: ofv216.

8. Chartrand C, Leeflang MMG, Minion J, Brewer T, Pai M. 2012. Accuracy of rapid influenza diagnostic tests: a meta-analysis. Ann Intern Med 156: 500–511.

9. Chartrand C, Tremblay N, Renaud C, Papenburg J. 2015. Diagnostic accuracy of rapid antigen detection tests for respiratory syncytial virus infection: systematic review and meta-analysis. J Clin Microbiol 53: 3738–3749.

10. Merckx J, Wali R, Schiller I, Caya C, Gore GC, Chartrand C, Dendukuri N, Papenburg J. 2017. Diagnostic accuracy of novel and traditional rapid tests for influenza infection compared with reverse transcriptase polymerase chain reaction: a systematic review and meta-analysis. Ann Intern Med 167: 394–409.

11. Banerjee D, Kanwar N, Hassan F, Essmyer C, Selvarangan R. 2018. Comparison of six sample-to-answer influenza A/B and respiratory syncytial virus nucleic acid amplification assays using respiratory specimens from children. J Clin Microbiol 56: e00930–18.

12. Banerjee D, Kanwar N, Hassan F, Lankachandra K, Selvarangan R. 2019. Comparative analysis of four sample-to-answer influenza A/B and RSV nucleic acid amplification assays using adult respiratory specimens. J Clin Virol 118: 9–13.

13. Binnicker MJ, Espy MJ, Irish CL, Vetter EA. 2015. Direct detection of influenza A and B viruses in less than 20 minutes using a commercially available rapid PCR assay. J Clin Microbiol 53: 2353–2354.

14. de-Paris F, Beck C, Machado AB, Paiva RM, da Silva Menezes D, de Souza Nunes L, Kuchenbecker R, Barth AL. 2012. Optimization of one-step duplex real-time RT-PCR for detection of influenza and respiratory syncytial virus in nasopharyngeal aspirates. J Virol Methods 186: 189–192.

15. Focus Diagnostics Inc. 2016. Simplexa(tm) Flu A/B & RSV Package Insert, Cypress, CA, USA.

16. Kamau E, Agoti CN, Lewa CS, Oketch J, Owor BE, Otieno GP, Bett A, Cane PA, Nokes DJ. 2017. Recent sequence variation in probe binding site affected detection of respiratory syncytial virus group B by real-time RT-PCR. J Clin Virol 88: 21–25.

17. Melchers WJG, Kuijpers J, Sickler JJ, Rahamat-Langendoen J. 2017. Lab-in-a-tube: real-time molecular point-of-care diagnostics for influenza A and B using the cobas^®^ Liat^®^ system. J Med Virol 89: 1382–1386.

18. Nolte FS, Gauld L, Barrett SB. 2016. Direct comparison of Alere i and cobas Liat influenza A and B tests for rapid detection of influenza virus infection. J Clin Microbiol 54: 2763–2766.

19. Ling L, Kaplan SE, Lopez JC, Stiles J, Lu X, Tang Y-W. 2018. Parallel validation of three molecular devices for simultaneous detection and identification of influenza A and B and respiratory syncytial viruses. J Clin Microbiol 56: e01691–17.

20. Young S, Illescas P, Nicasio J, Sickler JJ. 2017. Diagnostic accuracy of the real-time PCR cobas^®^ Liat^®^ Influenza A/B assay and the Alere i Influenza A&B NEAR isothermal nucleic acid amplification assay for the detection of influenza using adult nasopharyngeal specimens. J Clin Virol 94: 86–90.

